# PCR-based amplification of a *cox1* mini-DNA barcode gene from feces: A non-invasive molecular technique to identify environmental DNA samples of maritime shrew (*Sorex maritimensis*)

**DOI:** 10.1101/2023.09.20.558484

**Authors:** Golnar Jalilvand, Donald T. Stewart

## Abstract

The Maritime Shrew (*Sorex maritimensis*) is endemic to Canada and found only in Nova Scotia and New Brunswick. The Maritime Shrew has been identified as one of the vertebrate species in Nova Scotia that is most susceptible to the effects of climate change and global warming, and it is listed by NatureServe as Vulnerable (category G3). While generally regarded as a wetland specialist, relatively little is known about their specific habitat preferences. Non-invasive methods of sampling have proven valuable in identifying and monitoring such rare species. The objective of this study was to optimize a non-invasive method to document presence of Maritime Shrews using non-invasively collected fecal DNA and to develop a PCR-based protocol to amplify a short, ∼120 base pair section of the *cox1* gene using shrew-specific primers. We used baited feeding tubes to collect shrew feces. C*ox1* PCR primers were designed to preferentially amplify this mini-DNA barcode for shrews in samples that may contain feces from rodents as well. The primers were designed to amplify a small amplicon to increase the likelihood of successful amplification from degraded DNA. This technique is likely to be effective for documenting the distribution and habitat preferences of this relatively rare shrew in Nova Scotia and New Brunswick.

## Introduction

Genetic material is transferred from organisms to their environment in various ways, such as through secretions, blood, scat, urine, etc. (Thomsen and Willerslev 2015). Animal scats and regurgitated pellets are a possible source of DNA in the environment (Bowles et al. 2011). Genetic analysis of these materials can be used to monitor the occurrence, spread and/or population decline of small mammals, and it is a valuable tool for researchers and wildlife conservationists (McInnes et al. 2017, Murray et al. 2011, O’Meara 2014). Environmental DNA (eDNA) analysis is an effective method to detect species across a wide array of aquatic and terrestrial ecosystems. This method is an improvement on traditional sampling methods such as kill sampling or catch and release trapping in terms of sampling effort and sensitivity to detection of the targeted species (Deiner et al. 2017, Dejean et al. 2012, Vynne et al. 2012). Environmental DNA degradation, however, can limit the application of eDNA analysis, especially in warm habitats, where only small segments of DNA are likely to remain intact for more than a day or two (Deiner et al. 2017, Goldberg et al. 2016, Hering et al. 2018). Amplification success rates are higher from fresh droppings or scats that are deposited dry or that quickly dry out (McInnes et al. 2017). Moreover, scat samples may include non-target DNA originating from parasites in animal scat, gut flora, and their food;droppings may become contaminated by external organisms such as insects and vegetation in the environment or from material from other mammals that co-occurred at the same location (McInnes et al. 2017). Given the challenges of analysis of DNA from certain types of degraded samples, it is helpful to design primers to amplify specific, short regions of the DNA barcoding *cox1* gene of a target species that can be used to detect, identify, and distinguish the target species’ DNA easily and efficiently (Meusnier et al. 2008).

The Maritime Shrew (*Sorex maritimensis* Smith) is a Canadian endemic with a limited distribution in Nova Scotia and New Brunswick and it is the only mammal endemic to that region (Dawe et al. 2009, McAlpine et al. 2012, Stewart et al. 2002). The Maritime Shrew is primarily associated with grass–sedge marshes, low-lying floodplains, wet meadows, and marsh margins (Dawe 2004, Herman and Scott 1994, McAlpine et al. 2012, Perry et al. 2004, Scott and Hebda 2004). More specifically, Dawe (2004) identified an association of the Maritime Shrew with abundant graminoids (i.e., grasses, sedges, and rushes), particularly *Calamagrostis canadensis* (Michx.) (Bluejoint Reedgrass), and low tree cover. These habitats constitute a somewhat limited and fragmented habitat at the terrestrial/aquatic ecotone in the Maritimes, which contributed to the Maritime Shrew being ranked as the most vulnerable mammal in Nova Scotia to the effects of climate change and global warming (Herman and Scott 1992). Various small mammal surveys have expanded the known habitats where Maritime Shrews can be found. For example, McAlpine et al. (2012) collected one Maritime Shrew in northern New Brunswick in habitat characterized as a marshy-meadow located within a mixed-forest of *Thuja occidentalis* L. (Eastern White Cedar), *Abies balsamia* (L.)(Balsam Fir), *Picea rubens* Sarg. (Red Spruce), and *P. mariana* (Mill.) (Black Spruce). Henderson and Forbes (2012) similarly documented presence of Maritime Shrews in what was previously considered atypical habitat for the species, namely, a predominantly Black Spruce forest with large amounts of *Pinus strobus* L. (Eastern White Pine), and some *Acer rubrum* L. (Red Maple), *Betula* spp. L. (birches), and *Alnus* spp. L. (alders).

As suggested by Henderson and Forbes (2012), in light of additional information on habitat associations of the Maritime Shrew, more precise ecological data are needed to predict the distribution of suitable habitats in northeastern North America to propose management plans to conserve this Canadian endemic mammal. Meanwhile, improvement in identification techniques of small mammal feces can be a valuable tool for detecting the presence of rare and endangered species and obtaining further data on habitat preferences of this species. In this study, we present a PCR-based protocol to identify the Maritime Shrew from feces collected using the baited feeding tube method developed by Churchfield and colleagues (Churchfield et al. 2000, Greenwood et al. 2002).

### Field-site description

Based on the apparent preference of the Maritime Shrew for wetlands (Dawe 2004, Smith 1940), the field-site for collecting feces for this project was the Wolfville dykelands, bordering the Minas Basin, Nova Scotia. This is also the area where the type specimen of the species was first collected by Smith (1940). The specific field-site is located on artificial dykelands first created by Acadian settlers in this region of Nova Scotia (see Bleakney 2004) so the land is below sea-level and generally quite moist. The dykelands consist of three distinct zones: (1) the dykes themselves, which create (2) areas inside the dykes,which are referred to here as the “protected dykelands”, and (3) the seaward, tidal, wetland area dominated by *Juncus* spp. (rushes) and *Spartina* spp. (cordgrasses) (Hatcher and Patriquin 1981). The dykes were constructed to increase agricultural lands, and this continues to be how the dykelands are generally used today (Sherren et al. 2021). The parts of the protected dykelands that are not maintained for agriculture are characterized by old-field vegetation including perennial grasses and forbs such as Goldenrod (*Solidago* spp.), meadowsweet (*Spirea latifolia* W. Aiton), Canada blue-joint grass (*Calamagrostis canadensis* (Michx.) P. Beauv.) and sedges (*Carex* spp.), as well as wild rose (*Rosa virginiana* Mill.).

## Methods

### Collection of fecal samples and extraction of DNA

All methods were approved by the Acadia University Animal Care Committee, and met guidelines set by the Canadian Council on Animal Care. Fieldwork opportunities for this project were interrupted by the COVID-19 pandemic. Two brief field sessions (November 2019 and September 2020) allowed collection of sufficient material to test the effectiveness of the feeding tube method for collecting feces and the newly developed shrew specific mini-DNA barcode primers. Forty feeding tubes (Greenwood et al. 2002) were constructed from 5 cm wide white plastic PVC pipe cut to lengths of 25 cm. Each tube was covered on one end with fiberglass mesh window screen (Climaloc™) and secured with duct tape (Fig, 1). A piece of white parchment paper 20 x 10 cm was placed in each tube to make scat collection easier. Bait consisted of 2 or 3 frozen mealworm beetle larva placed near the mesh end of each tube. Feeding tubes were positioned haphazardly between grasses and herbaceous plants and were checked twice a day, shortly after sunrise and just before sunset. Immediately after collection, scat samples were transferred to a -40°C freezer to limit further DNA degradation. Feeding tubes were washed with dilute Alconox™ detergent in tap water, rinsed thoroughly in tap water, rinsed again in reverse osmosis distilled water, and then air-dried before being returned to the field. A Quick-DNA Fecal/Soil Microbe 96 Kit (Zymogen) was used for DNA isolation of the fecal samples according to the protocol provided with the kit. The amount of fecal material used per isolation typically ranged from 50 to 150 mg of material collected from a single feeding tube. The concentration and purity of isolated DNA were quantified using a BioDrop DUO+ Micro-volume Spectrophotometer. A summary of the workflow is shown in Figure 2.

**Figure 1.**
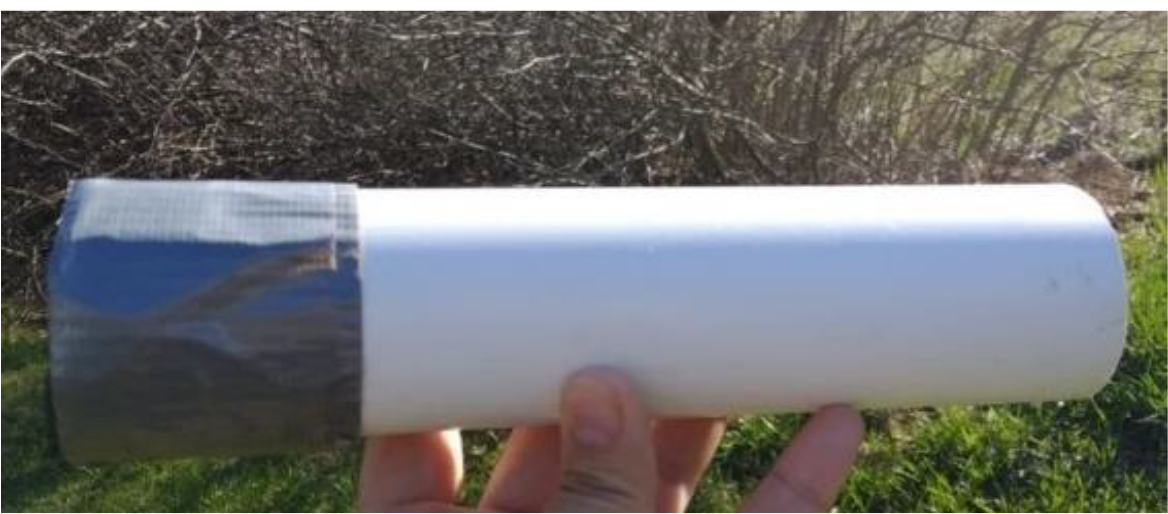
Baited feeding tubes made from 25 cm lengths of PVC plastic pipe. Photograph © XXXX.

**Figure 2.**
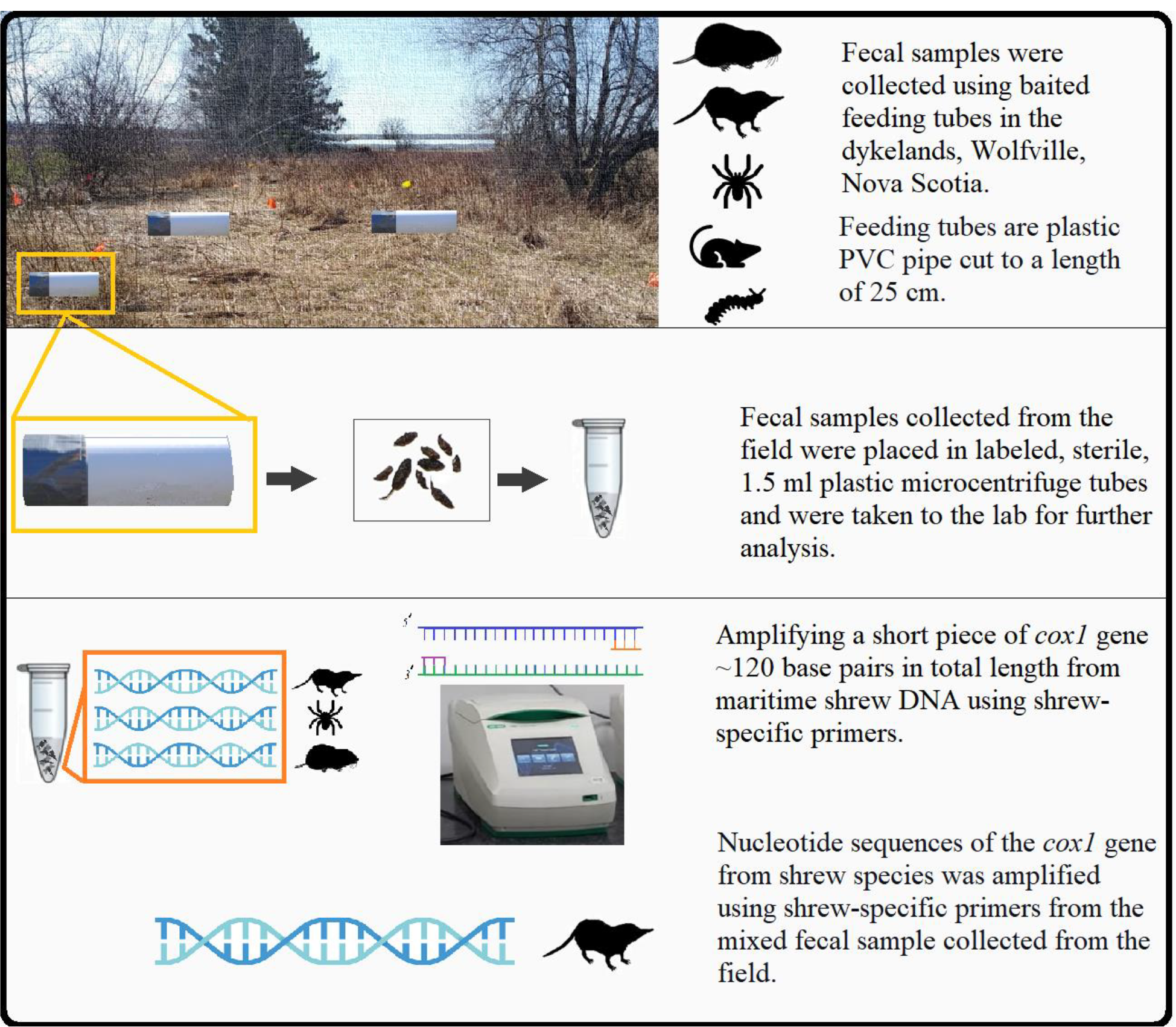
Summary of the work-flow. Photograph © XXXX.

### Design of PCR primers and PCR conditions

A set of forward and reverse test primers were designed using Primer3 (https://bioinfo.ut.ee/primer3-0.4.0/). A 657-bp fragment of the mitochondrial *cox1* gene from a specimen of *Sorex maritimensis* (recognized as *Sorex arcticus* [Arctic Shrew] on GenBank, accession number JF436020) from Windsor, Nova Scotia, was input into Primer3. We specified amplification ofa short, ∼120bp fragment of the *cox1* gene from Maritime Shrew to increase the probability of successful amplification from degraded fecal DNA. It was initially not known whether this fragment might also amplify *cox1* from other shrew species. The resulting mini-barcoding primers were as follows: F221COI 5’-CAGACATAGCATTCCCACGA-3’ and R345COI 5’-TGGGGGATAGACAGTTCAGC-3’. Forward and reverse M13 primer sequence tags were added to the forward and reverse *cox1* primer sequences, respectively, to facilitate Sanger sequencing at the Genome Quebec sequencing facility at McGill University in Montréal (see https://cesgq.com/en-faq).

Double-stranded amplification of the *cox1* mini-barcoding fragment using primers F221COI and R345COI was performed in 50 µl reactions using the following cycling parameters: initial denaturation at 94°C for 4 min, followed by 39 cycles of 30 seconds denaturation at 94°C, 60 seconds annealing at 55°C, and 30 seconds extension at 72°C. The cycling period was followed by an additional 72°C extension step of 5 minutes after which the reactions were paused at 5°C. Reactions contained 25 µl BIO-RAD Master Mix, 3 µL each forward and reverse primers (10 mM), 1–2 µL DNA template, and 17–18 µL ultrapure water. Successfully amplified products were sent to the McGill University and Génome Québec Innovation Centre for Sanger sequencing. The CAP3 Sequence Assembly Program (http://doua.prabi.fr/software/cap3) was used to merge forward and reverse sequences. Sequences were analyzed with MEGA 7.0.26 (Kumar et al., 2016).

### Identification of shrew species

For species identification, FASTA formatted sequences were submitted to the NCBI BLAST site (https://blast.ncbi.nlm.nih.gov/Blast.cgi); “Highly similar sequences” was selected, and the Organism option was set to “Soricidae (taxid:9376)”.

### Identification of non-target species

It was likely that some fecal samples belonged to other species, including rodents, in which case amplification with shrew specific primers would not result in PCR products. To test this hypothesis, four of the fecal samples that gave no PCR product with the shrew specific primers were amplified with universal *cytochrome b* primers L14841 5’-AAAAAGCTTCCATCCAACATCTCAGCATGATGAAA and H15149 5’-AAACTGCAGCCCCTCAGAATGATATTTGTCCTCA (Irwin et al. 1991, Kocher et al. 1989), which should amplify a 307 bp fragment of the mammalian cyt *b* gene.

## Results and Discussion

In total, forty-three sets of fresh small mammal scats were collected using 40 feeding tubes in two field sessions, one session in November 2019 and the second session in September 2020 “A”. Twenty-seven fecal samples were collected during the first sampling period and sixteen fecal samples during the second period. A total of seventeen sets of fresh scats that looked like shrew feces (i.e., they were ∼3–4 mm in length, slightly twisted/coiled with tapered ends) were selected for further analysis (Chame 2003).

From the DNA isolated from these scats, 6/17 (35.29%) successfully amplified with the shrew-specific primers. Of these 6 samples, 5 matched *S. maritimensis* on GenBank. Although the samples are listed as Arctic Shrews on GenBank, Arctic Shrews from Nova Scotia and New Brunswick, which were traditionally classified as *S. arcticus maritimensis*, have been reclassified as *S. maritimensis* (Stewart et al. 2002, Shafer and Stewart 2007). The sixth sample was a close match to a Masked Shrew (*S. cinereus* Kerr) (GenBank Accession number JF436364.1) from Nova Scotia. Several samples amplified with the universal mammalian primers L14841 and H15149; however, not all of these returned usable sequence. Two samples were highly similar (99.0%) to Meadow Vole (*Microtus pennsylvanicus* [Ord 1815]), and one was highly similar (98.2%) to White-footed mouse (*Peromyscus leucopus* [Rafinesque 1818]). The Meadow Vole is common in grassland habitats throughout Nova Scotia (Scott and Hebda 2004).

In conclusion, these findings demonstrate that this primer set can be applied to studies of the habitat use and preferences of the Maritime Shrew, *Sorex maritimensis*. Application of this technique throughout Nova Scotia, New Brunswick, and other regions where the Maritime Shrew has not been previously documented such as Prince Edward Island, Canada and Maine, USA, will help clarify the distributional range of this species.

